# Uncovering the role of admixture in disease and drug response: Association of hepatocyte gene expression and DNA methylation with African Ancestry in African Americans

**DOI:** 10.1101/491225

**Authors:** CS Park, T De, Y Xu, Y Zhong, ER Gamazon, E Smithberger, C Alarcon, MA Perera

## Abstract

**Background:** African Americans (AAs) are an admixed population with portions of their genome derived from West Africans and Europeans. In AAs, the proportion of West African ancestry (WAA) can vary widely and may explain the genetic drivers of disease, specifically those that disproportionately affect this understudied population. To examine the relationship between the proportion of WAA and gene expression, we used high dimensional data obtained from AA primary hepatocytes, a tissue important in disease and drug response.

**Methods:** RNA sequencing (Illumina HiSeq Platform) was conducted on 60 AA-derived primary hepatocytes, with methylation profiling (Illumina MethylationEPIC BeadChip) of 44 overlapping samples. WAA for each sample was calculated using fastSTRUCTURE and correlated to both gene expression and DNA methylation. The GTEx consortium (n = 15) was used for replication and a second cohort (n = 206) was using used for validation using differential gene expression between AAs and European-Americans.

**Results:** We identified 131 genes associated with WAA (FDR< 0.1), of which 28 gene expression traits were replicated (FDR<0.1) and enriched in angiogenesis and inflammatory pathways (FDR<0.1). These 28 replicated gene expression traits represented 257 GWAS catalog phenotypes. Among the PharmGKB pharmacogenes, *VDR, PTGIS, ALDH1A1, CYP2C19* and *P2RY1* were associated with WAA (*p* < 0.05) with replication of *CYP2C19* and *VDR* in GTEx. Association of DNA methylation with WAA identified 1037 differentially methylated regions (FDR<0.05), with hypomethylated genes enriched in drug response pathways. Overlapping of differentially methylated regions with the 131 significantly correlated gene expression traits identified 5 genes with concordant directions of effect: *COL26A1, HIC1, MKNK2, RNF135, SNAI1* and *TRIM39*.

**Conclusions:** We conclude that WAA contributes to variability in hepatic gene expression and DNA methylation with identified genes indicative of diseases disproportionately affecting AAs. Specifically, WAA-associated genes were linked to previously identified loci in cardiovascular disease (*PTGIS, PLAT*), renal disease (*APOL1*) and drug response (*CYP2C19*).

## Background

African Americans (AAs) are an admixed population, having varying proportions of African and European ancestry between individuals [1]. As a consequence of their West African ancestry (WAA), AAs have more genetic variation and shorter extent of linkage disequilibrium (LD) than European Ancestry populations, with the proportion of WAA varying greatly between self-identified AAs [2]. It is this genetically driven variability that may aid in explaining differences in hepatic gene expression and DNA methylation patterns, which cannot be elucidated in homogenous populations such as those of European-only ancestry. For example, WAA has been shown to predict a stronger inflammatory response to pathogens compared to those of European ancestry due to recent selective pressures specific to this population [3].

Furthermore, AAs suffer disproportionately from many chronic diseases and adverse drug reactions, as compared to other populations [4, 5] as well as are protected from some conditions. AAs have a higher risk of cardiovascular events and negative outcomes therapy [6]. They have higher incidences of death and disability from cardiovascular diseases (CVDs), thrombosis, renal dysfunction and pathologies, diabetes, cancers, and other metabolic disorders [7-15]. Conversely, they have lower prevalence of disease such as testicular cancer [16]. Differences in gene expression may help explain these observed differences.

Due to the key role of the liver in biosynthesis, drug metabolism and complex human diseases, genetic and epigenetic differences in the liver may uncover the underlying causal genes responsible for chronic diseases that disproportionately affect AAs [17, 18]. The first comprehensive mapping of liver expression quantitative trait loci (eQTLs) proposed several candidate susceptibility genes associated with type I diabetes, coronary artery disease and plasma cholesterol levels in a white cohort [19]. More recently, finer-resolution mapping of liver eQTLs, combining both gene expression data with histone modification-based annotation of putative regulatory elements, identified 77 loci found to associate with at least one complex phenotype [20]. In addition, our group has previously shown that studies specifically investigating the unique genetic variants of AAs can reveal population-specific risk factors that may explain differences in drug response, such as African ancestry-specific genetic risk factors associated to a higher risk of bleeding from warfarin therapy in AAs [12] as well as population specific variants associated with increased risk of thrombotic disease [21].

Rather than identifying disease susceptibilities through genome-wide association studies (GWAS), here we use variability in genetic ancestry to uncover potential drivers of disease and drug response that may potentially explain differences in disease and drug response in AAs. In this study we correlated WAA to both gene expression and differentially methylated regions to determine the contribution of WAA admixture in hepatic gene expression traits. We identify genes are known to be dysregulated in diseases, which affect AAs disproportionately and may also be responsible for adverse drug responses in AA populations.

## Materials and methods

### Cohorts

A total of 68 African American (AA) primary hepatocyte cultures were used for this study. Cells were either purchased from commercial companies (BioIVT, TRL, Life technologies, Corning and Xenotech), or isolated from cadaveric livers using a modified two-step collagenase perfusion procedure [77]. Liver specimens were obtained through collaborations with Gift of Hope, which supplies non-transplantable organs to researchers. In addition, we used GWAS data for 153 subjects from Genotype-Tissue Expression Project (GTEx) release version 7, of which 15 samples were used as a replication set.

### Primary Hepatocyte Isolation

Briefly, cadaveric livers obtained from Gift of Hope were transferred to the perfusion vessel Büchner funnel (Carl Roth) and the edge was carefully examined to locate the various vein and artery entries that were used for perfusion. Curved irrigation cannulae with olive tips (Kent Scientific) were inserted into the larger blood vessels on the cut surface of the liver piece. The liver was washed by perfusion of 1 L Solution 1 (HEPES buffer, Sigma-Aldrich), flow rate 100-400 mL/min, with no recirculation, followed by perfusion with 1 L of Solution 2 (EGTA buffer, Sigma-Aldrich), flow rate 100-400 mL/min, with no recirculation. The tissue was washed to remove the EGTA compound by perfusion of 1 L Solution 1, flow rate 100-400 mL/min, with no recirculation. The liver was digested by perfusion with Solution 3 (collagenase buffer, Sigma-Aldrich), flow rate 100-400mL/min, with recirculation. Following perfusion, liver section was placed in a crystallizing dish (Omnilab) containing 100-200 ml of Solution 4 (Bovine Serum Albumin, Sigma-Aldrich). The Glisson’s capsule was carefully removed and the tissue was gently shaken to release hepatocytes. The cell suspension was then filtered by a 70 μm nylon mesh (Fisher Scientific), and centrifuged at 72 x g for 5 min at 4 °C. The pellets contained hepatocytes that was washed twice with solution 4 and resuspended in plating medium (Fisher Scientific).

For primary hepatocyte cultures, cell viability was determined by trypan blue (Lonza) exclusion using a hemocytometer (Fisher Scientific) [78]. If viability was low, Percoll gradient (Sigma-Aldrich) centrifugation of cell suspensions was carried out to improve yield of viable cells. Cell were plated at a density of 0.6 × 10^6^ cells/well in 500 μL InVitroGro-CP media (BioIVT, Baltimore, MD) in collagen-coated plates with matrigel (Corning, Bedford, MA) overlay and incubated overnight at 37° C. Cultures were maintained in InVitroGro-HI media (BioIVT) supplemented with Torpedo antibiotic mix (BioIVT) per the manufacturer’s instructions. The media was replaced every 24 hours for three days. RNA was extracted after three days using the RNAeasy Plus mini-kit (Qiagen) per manufacturer’s instructions.

### Genotyping and quality control

DNA was extracted from each hepatocyte line using Gentra Puregene Blood kit (Qiagen) as per manufacturer’s instructions from 1-2 million cells. All DNA samples were then bar-coded for genotyping. SNP genotyping was conducted using the Illumina Multi-Ethnic Genotyping array (MEGA) at the University of Chicago’s Functional Genomics Core using standard protocols.

Quality control (QC) steps were applied as previously described including an w imputation info metric threshold of > 0.4 [21]. Briefly, a sex check was performed on genotypes using PLINK (version 1.9) to identify individuals with discordant sex information. Duplicated or related individuals were identified using identity-by-descent (IBD) method with a cutoff score of 0.125 indicating third-degree relatedness. A total of five individuals were excluded after genotyping QC analysis. SNPs located on the X and Y chromosomes and mitochondrial SNPs were excluded. SNPs with a missing rate of > 5% or those that failed Hardy-Weinberg equilibrium (HWE) tests (*p* < .00001) were also excluded.

### African ancestry measurement

The genotypes of 68 primary hepatocytes and 153 GTEx subjects were merged with HapMap phase 3 reference data from four global populations; Yoruba in Ibadan, Nigeria (YRI); Utah residents with Northern and Western European ancestry (CEU); Han Chinese in Beijing, China (CHB); and Japanese in Tokyo, Japan (JPT) [79]. Population structure of the merged data was inferred by the Bayesian clustering algorithm STRUCTURE deployed within fastStructure v1.0 and performed without any prior population assignment. We employed the admixture model, and the burn in period and number of Markov Chain Monte Carlo repetition was set to 20,000 and 100,000, respectively [80]. The number of parental populations (K) was set to 3, purporting three main continental groups (African, European, or Asian). WAA percentages of the primary hepatocytes and GTEx subjects were calculated as the probability of being grouped as Yoruba African, Caucasian, and East Asian, respectively [80]. All individuals in our 60 AA cohort had WAA greater than 40%.

### GTEx replication liver cohort

Of the 144 GTEx liver IDs extracted from “GTEX_Sample_Attributes” file, five were replicates and were removed from the analysis. Among the remaining 139 unique liver IDs, 118 had available genotype information for ancestry determination. Individuals with WAA greater than 40% were included in the GTEx AA Liver cohort. After WAA estimation, normalized gene expression reads for 15 subjects meeting the ancestry inclusion criteria were extracted from GTEx expression file (GTEx_Liver.v7.normalized_expression.bed). Age and sex information of these subjects were extracted from the subject phenotype file on the GTEx Portal site (GTEx_v7_Annotations_SubjectPhenotypesDS.txt).

### Validation analysis in an independent liver transcriptome dataset

Gene expression profiling in liver had previously been conducted in 206 samples using Agilent-014850 4×44 k arrays (GPL4133) [23, 81]. These samples had come from donor livers not intended for organ transplantation. Genotyping on these samples had been done using the Illumina Human 610 quad beadchip platform (GPL8887) at the Northwestern University Center for Genetic Medicine Genomics Core Facility and imputation was subsequently performed using bimbam [82]. Principal component analysis was used to quantify ancestry using the Human Genome Diversity Panel with African and European populations as reference, as previously described [23].

We conducted differential expression analysis between the European samples and the African American samples using Linear Models for Microarray Data (*limma*) as implemented in the Bioconductor package [83]. This Bayesian methodology uses a “moderated” t-statistic from the posterior variance assuming an inverse chi-squared prior for the unknown variance for a gene. We used Bonferroni-adjusted p<0.05 based on the total number of genes that were tested for replication.

### RNA-sequencing and quality control

Total RNA was extracted from each primary hepatocyte culture after three days in culture using the Qiagen Rneasy Plus mini-kit per manufacturer’s instructions. RNA-QC was performed using an Agilent Bio-analyzer and samples with RNA integrity number (RIN) scores above 8 were used in subsequent sequencing. RNA-Seq libraries were prepared for sequencing using Illumina mRNA TruSeq RNA Sample Prep Kit, Set A (Illumina catalog # FC-122-1001) according to manufacturer’s instructions. The cDNA libraries were prepared and sequenced using both Illumina HiSeq 2500 and HiSeq 4000 machines by the University of Chicago’s Functional Genomics Core to produce single-end 50 bp reads with approximately 50 million reads per sample (accession number GSE124076). Batch effects were corrected for in quality control below.

Quality of the raw reads was assessed by FastQC (version 0.11.2). Fastq files with a per base sequence quality score > 20 across all bases were included in downstream analysis. Reads were aligned to human Genome sequence GRCh38 and comprehensive gene annotation (GENCODE version 25) was performed using STAR 2.5. Only uniquely mapped reads were retained and indexed by SAMTools (version 1.2). Nucleotide composition bias, GC content distribution and coverage skewness of the mapped reads were further assessed using read_NVC.py, read_GC.py and geneBody_coverage.py scripts, respectively, from RseQC (version 2.6.4). Samples without nucleotide composition bias or coverage skewness and with normally distributed GC content were reserved. Lastly, Picard CollectRnaSeqMetrics (version 2.1.1) was applied to evaluate the distribution of bases within transcripts. Fractions of nucleotides within specific genomic regions were measured for QC and samples with > 80% of bases aligned to exons and UTRs regions were considered for subsequent analysis.

### RNA-seq data analysis

Post alignment and QC, reads were mapped to genes referenced with comprehensive gene annotation (GENCODE version 25) by HTSeq (version 0.6.1p1) with union mode and minaqual = 20 [84]. HTSeq raw counts were supplied for gene expression analysis using Bioconductor package DESeq2 (version1.20.0) [85]. Counts were normalized using regularized log transformation and principal component analysis (PCA) was performed in DESeq2. PC1 and PC2 were plotted to visualize samples expression pattern. Three samples with distinct expression patterns were excluded as outliers resulting in 60 samples used in RNA-seq analysis. We calculated TPM (transcript per million) by first normalizing the counts by gene length and then normalizing by read depth [86]. Gene expression values were filtered based on the expression thresholds > 0.1 TPM in at least 20% of samples and ≥ 6 reads in at least 20% of samples as done in the GTEx consortium (gtexportal.org, Analysis Methods, V7, updated 09/05/2017).

WAA percentage, gender, age, platform and batch were used as covariates for downstream analysis. Probabilistic estimation of expression residuals (PEER) method v1.3 was used to identify PEER factors and linear regression was run on inverse normal transformed expression data using five PEER factors, based on GTEx’s determination of number of factors as a function of sample size [88, 89]. WAA-associated genes were identified from a genome-wide list of 18,854 genes for our hepatocyte-derived AA cohort samples and replicated in 21,730 genes from the GTEx-derived AA cohort at FDR cutoff of 0.10. The top 131 genes with an FDR < 0.10 were also replicated in the independent Replication cohort obtained from Innocenti et al. (GEO accession number GSE26106) [23]. The 28 replicated genes were used to identify overlap with the NHGRI-EBI GWAS catalog reported genes to identify GWAS traits link to our replicated genes, v1.0.2 [24]. FDR calculations from linear regressions on gene expression were conducted with the “p.adjust” function in R and the default method of “fdr” was used.

In addition, we also conducted a subset analysis with 64 PharmGKB “very important pharmacogenes”. These genes have extensive literature support for association to drug responses. We analyzed this subset for association with WAA using the same linear regression method at a nominal *p*-value cutoff of 0.05 due to the smaller number of genes being tested, 64, our smaller sample sizes of 60 in our AA cohort and 15 in the GTEx replication cohort.

### Methylation sample preparation and data analysis

DNA was isolated from hepatocytes or liver tissue. Liver tissue was homogenized in a bead mill (Fisher Scientific) using 2.8 mm ceramic beads. Then, DNA from hepatocytes or liver was extracted with the Gentra Puregene Blood kit (Qiagen) as per manufacturer instructions. One microgram of DNA was bisulfite converted at the University of Chicago Functional Genomics Core using standard protocols. CT-conversion was performed using Zymo-Research EZ DNA Methylation kits and further processed for array hybridization using Illumina provided array reagents. Following hybridization the arrays were stained pre manufacturer’s protocol and analysed using an Illumina HiSCAN. Of the 60 available hepatocyte cultures only 56 produced sufficient bisulfite converted DNA for analysis.

Illumina MethylationEPIC BeadChip microarray (San Diego, California, USA), consisting of approximately 850,000 probes, predefined and annotated [90], and containing 90% of CpGs on the HumanMethylation450 chip and with more than 350,000 CpGs regions identified as potential enhancers by FANTOM5 [91] and the ENCODE project [92], was used for methylation profiling of DNA extracted from 56 AA hepatocytes, which overlapped the samples used for gene expression analysis (accession number GSE123995). Raw probe data was analyzed using the ChAMP R package for loading and base workflow [26], which included the following R packages: BMIQ for type-2 probe correction method [93]; ComBat for correction of multiple batch effects including Sentrix ID, gender, age, slide and array [94, 95]; svd for singular value decomposition analysis after correction [96]; limma to calculate differential methylation using linear regression on each CpG site with WAA as a numeric, continuous variable [97]; DMRcate for identification of DMRs, using default parameters, and the corresponding number of CpGs, minimum FDR (minFDR); Stouffer scores, and maximum and mean Beta fold change values [98]; minfi for loading and normalization [99]; missMethyl for gometh function for GSEA analysis [100]; and FEM for detecting differentially methylated gene modules [28].

Methylation data quality control in ChAMP’s *champ.load()* and *champ.filter()* function removed the following probes: 9204 probes for any sample which did not have a detection *p-*value < 0.01, and thus considered as a failed probe, 1043 probes with a bead count < 3 in at least 5% of samples, 2975 probes with no CG start sites, 78,753 probes containing SNPs [101], 49 probes that align to multiple locations as identified in by Nordlund et al. [102], and 17,235 probes located on X and Y chromosomes. Threshold for significantly differentially methylated probes was set at an adjusted *p*-value of 0.05 using Benjamini-Hochberg correction for multiple testing in the limma package. Significance for DMRs was set at adjusted FDR < 0.05 in the DMRcate package using default parameters. Analysis was performed with R statistical software (version 3.4.3 and version 3.5 for ChAMP (version 2.10.1)). Three outliers and nine samples from young subjects, less than five years old were excluded due to known differences in methylation profiles associated with age [103] leaving 44 samples in the analysis.

Gene ontology analysis was performed using g:Profiler (biit.cs.ut.ee/gprofiler) using g:GOSt to provide statistical enrichment analysis of our significantly HypoM and HyperM genes, and significantly expressed genes associated with African ancestry [104]. We filtered for significance at a Benjamini-Hochberg adjusted *p*-value < 0.05, we set hierarchical sorting by dependencies between terms on the strongest setting, “best per parent group”, and only annotated genes were used as the statistical domain size parameter to determine hypergeometrical probability of enrichment. We considered ontology terms to be statistically significance at a BH-adjusted *p*-value < 0.01.

Correlation of log fold change in DNA methylation of DM CpG sites by genomic feature was performed on significantly DM probes identified by limma and plotted using ggplot2 v3.1.0. Genomic feature and transcriptionally-regulated regions were subset for terms “island”, “opensea”, “shelf”, “shore” and “5’UTR”, “TSS1500”, “Body”, “3’UTR”, respectively. A chi square test was used for all categorical comparisons. Correlation with methylation and gene expression was performed using the ggscatter function in ggpubr v0.2, with the correlation method set to Pearson, confidence interval set at 95% and regression calculations included for the following subsets: HypoM, HyperM and cumulative CpG sites. Gene name conversions and annotated gene attributes were determined by BioMart v2.38.0. The Circos plot was created using Circos tool v0.69-6.

## Results

### Cohorts

60 primary hepatocyte cultures, procured from self-identified AAs, passed all quality control steps and were used for RNA-sequencing (RNA-seq) analysis. 56 hepatocyte cultures were assayed for DNA methylation, with 44 used in the final methylation analysis. We obtained genome-wide genotyping data of 153 subjects from GTEx liver cohort, version 7 [22]. All 60 of our AA hepatocyte cohort were confirmed as having WAA ranging from 41.8% to 93.7%, and 15 subjects in the GTEx liver cohort met the WAA inclusion criteria with WAA ranging from 75.6% to 99.9% (**Supplementary Figure 1**). Table 1 shows the demographics of each cohort. In general, the GTEx and the replication cohorts were older and had a lower percentage of females than the primary hepatocyte cohort.

**Table 1.** Demographics and clinical characteristics of hepatocyte/liver cohorts.

| Variable | RNA-Seq<br>Cohort | Methylation<br>Cohort | GTEx<br>Liver Cohort | Validation<br>Liver Cohort |
| --- | --- | --- | --- | --- |
|  | AA<br>(n = 60) | AA<br>(n = 44) | AA<br>(n = 15) | AA and EA<br>(n = 206) |
| Percent AA subjects<br>(%) | 100 | 100 | 100 | 11.1 |
| Age, years<br>(mean ± SD) | 39 ± 20.5 | 46 ± 12.2 | 52 ± 11.6 | 46 ± 22 |
| Sex (Female %) | 48.3 | 52.3 | 33.3 | 36.4 |
| West African<br>ancestry (%)<br>(mean ± SD) | 78.17 ± 12.86 | 79.83 ± 11.86 | 84.04 ± 7.13 | NA |

AA: African American greater than 40% YRI
EA: European American of CEU descent

**Figure 1.**
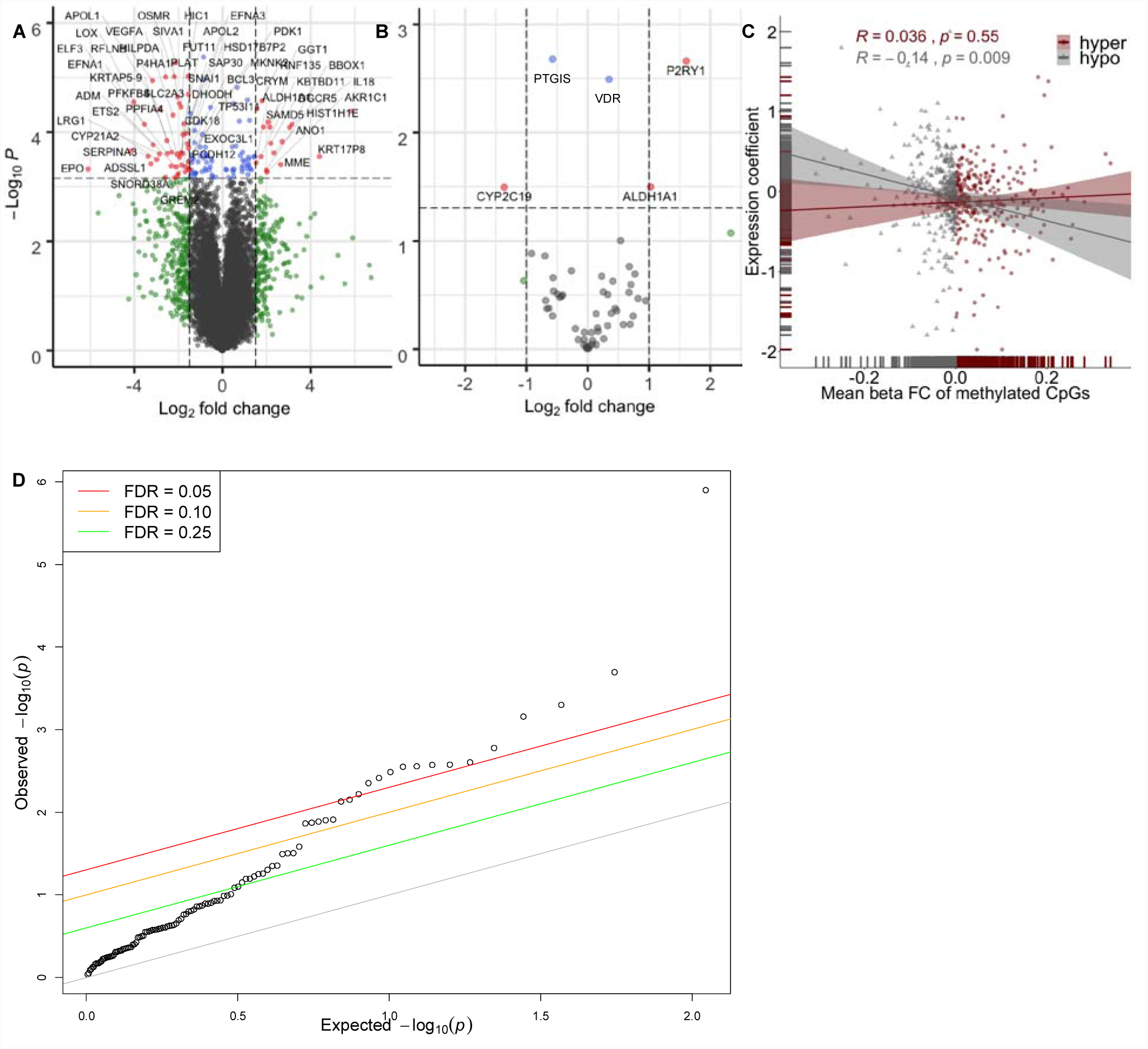
Gene expression traits and methylation patterns associated with West African ancestry in hepatocytes. Enhanced volcano plot of gene expression associated with increasing WAA plotted against – log_10_ *p*-values of **(A)** all 18,854 genes expressed in hepatocytes resulting in 131 genes significantly associated with WAA represented as red and blue dots (red circles: FDR < 0.05 and logFC > 1.5 and < −1.5; blue circles: FDR < 0.05 and logFC < 1.5 and > −1.5; green circles: FDR > 0.05 and logFC > 1.5 and < −1.5, grey circles: FDR > 0.05 and logFC < 1.5 and > −1.5) and **(B)** within subset of 64 PharmGKB “very important genes” (red circles: *p* < 0.05 and logFC > 1.0; blue circles: *p*-value < 0.05 and logFC < 1.0; grey circles: *p*-value > 0.05). **(C)** Correlation of 1034 unique genes containing DMRs significantly associated with WAA (mean Beta fold change from DMRcate) with coefficient of gene expression at each gene (indicating the direction of association to ancestry). Each point represents a gene, with grey triangles representing hypomethylated genes (Pearson’s r = −0.014, *p* = 0.009) and maroon red circles representing hypermethylated genes (Pearson’s r = 0.036, *p* = 0.55). Grey and maroon hash marks on the xand y-axis axis represent genes plotted with both expression and methylation values. Grey and maroon shading around each regression line represents the 95% confidence interval. **(D)** Q-Q plot of the observed versus expected – log_10_ *p*-values in the replication cohort (n = 206). Each point represents a gene with the colored lines representing different FDR thresholds of significance.

### Hepatic genes and pharmacogenes associated with African ancestry

Analysis of RNA-seq gene expression traits in the AA hepatocyte cohort with percentage WAA identified 131 genes, in which gene expression traits are significantly associated with WAA (**Supplementary Table 1**; **Figure 1A**, FDR < 0.10). We were able to replicate 28 of these genes in an independent dataset as differentially expressed (DE) between AAs and European-Americans (EAs) in liver [23] (**Table 2**; **Figure 1D**, *p* < 0.05). These 28 replicated gene expression traits are reported as trait-associated genes in 257 previous GWAS studies within the NCBI GWAS catalog [24]. These represent 119 unique traits (**Supplementary Table 2**) [24]. The phenotypic disease and biological GWAS traits associated with WAA include blood and blood pressure measures, coronary heart and artery disease, diabetic blood measures, chronic inflammatory disease, chronic kidney disease and various cancers.

**Table 2.** Significant and replicated DE genes between AAs and EAs (p < 0.05) from a genomewide discovery of genes associated with West African Ancestry (FDR < 0.10)

| Gene | AA Hepatocyte Cohort |  |  | DE in AA vs EA ‡ |
| --- | --- | --- | --- | --- |
| | Coefficient | $p$ | FDR < 0.10 | *$p < 0.05$ |
| **IL18** | 3.152 | 7.25E-05 | 0.0471 | 0.00000126 |
| **APOL1** | -2.848 | 3.83E-05 | 0.0380 | 0.000202 |
| **VEGFA** | -2.246 | 4.91E-05 | 0.0421 | 0.000502 |
| **PARD3** | 0.437 | 1.86E-04 | 0.0773 | 0.000696 |
| **SAP30** | -1.945 | 2.93E-04 | 0.0788 | 0.00167 |
| **LRRC37A2** | -0.306 | 6.73E-04 | 0.0988 | 0.00249 |
| **ENO1** | -1.007 | 1.89E-04 | 0.0773 | 0.00266 |
| **PGK1** | -0.929 | 5.29E-04 | 0.0882 | 0.00268 |
| **DGCR5** | 2.192 | 1.62E-04 | 0.0751 | 0.00278 |
| **MRO** | 1.131 | 2.57E-05 | 0.0380 | 0.00282 |
| **GREM2** | -2.191 | 6.86E-04 | 0.0988 | 0.00327 |
| **CYP21A2** | -2.986 | 3.17E-04 | 0.0790 | 0.00385 |
| **CEBPB** | -1.454 | 6.36E-04 | 0.0967 | 0.00445 |
| **MAD2L1BP** | -0.505 | 3.04E-04 | 0.0790 | 0.00605 |
| **GPR4** | -1.316 | 3.08E-04 | 0.0790 | 0.00706 |
| **RNF149** | -0.833 | 2.37E-04 | 0.0788 | 0.00743 |
| **GPI** | -1.453 | 4.58E-05 | 0.0411 | 0.0123 |
| **SLC22A15** | -1.271 | 6.86E-04 | 0.0988 | 0.0125 |
| **HIC1** | -1.567 | 9.64E-06 | 0.0307 | 0.013 |
| **APOL2** | -1.643 | 1.06E-04 | 0.0583 | 0.0134 |
| **MME** | 2.640 | 3.94E-04 | 0.0854 | 0.0137 |
| **MKNK2** | -1.545 | 3.44E-04 | 0.0811 | 0.0262 |
| **MGRN1** | -1.010 | 2.76E-04 | 0.0788 | 0.0313 |
| **C3orf33** | 0.506 | 6.45E-04 | 0.0973 | 0.0314 |
| **PTPN4** | 0.484 | 5.37E-04 | 0.0882 | 0.0321 |
| **NPR2** | 1.235 | 6.11E-05 | 0.0471 | 0.0444 |
| **MSX1** | 1.432 | 2.90E-04 | 0.0788 | 0.0448 |
| PDK1 | -2.038 | 6.30E-04 | 0.0966 | 0.0498 |

\* Replication $p$ -value ( $p < 0.05$ ) based on 131 discovery genes (Supplemental Table 1)
‡ Replication in differentially expressed (DE) genes between 23 African-Americans (AAs) and 183 European-Americans (EAs) [23]
Genes with FDR < 0.1 in the Replication Cohort are shown in **Bold**.

In the GTEx cohort of 15 AAs, we were able to replicate 8 of the 131 significant WAA-associated genes: *COL26A1* (effect size = −7.13, *p* = 0.037), *DHODH* (effect size = 3.19, *p* = 0.048), *GPI* (effect size = −3.46, *p* = 0.048), *HSD17B7P2* (effect size = −9.22, *p* = 0.027), *PLCL2* (effect size = −5.76, *p* = 0.046), *SLC2A3* (effect size = −5.95, *p* = 0.032), *TRIM39* (effect size = 5.53, *p* = 0.030) and *VEGFA* (effect size = −4.74, *p* = 0.032) in the GTEx liver cohort (**Supplementary Table 2**).

Due to the importance of the liver in pharmacologic drug response, we also tested the association of gene expression levels with percentage West African ancestry in a subset of genes belonging to the very important pharmacogenes (VIP) in PharmGKB, consisting of 64 genes known to be expressed in hepatocytes. These represent drug-metabolizing enzyme, transporters and drug target genes that are well-established for their role in drug response [25]. Testing the association between WAA and gene expression in the VIP genes identified five genes which were significantly associated to WAA: *VDR* (effect size = 0.35, *p* = 0.003), *PTGIS* (effect size = −0.57, *p* = 0.002), *ALDH1A1* (effect size = 1.03, *p* = 0.032), *CYP2C19* (effect size = −1.36, *p* = 0.032) and *P2RY1* (effect size = 1.61, *p* = 0.002 **(Figure 1B)**.

### DNA methylation patterns are associated with African ancestry in human liver

To identify differentially methylated (DM) regions (DMRs) and CpGs associated with WAA, we conducted linear regression on each CpG site to find African ancestry-related differential methylation [26]. We identified 23,317 significant DM CpG sites, out of a total of approximately 867,531 probes on the Illumina EPIC BeadChip microarray, annotated to 11,151 unique genes (**Supplementary Figure 2A**; BH-adjusted *p* < 0.05). These DM CpGs correspond to 1037 DMRs annotated to 1034 unique genes (**Supplementary Figure 2B**; minimum FDR < 0.0001). Each DM CpG site was categorized into hyper-(HyprM) and hypomethylated (HypoM) sites: 15,404 HyprM CpG sites constituted 435 HyprM DMRs, mapping to 432 unique genes, 7913 HypoM CpG sites constituted HypoM 602 DMRs, mapping to 602 unique genes, and seven annotated genes had both HyprM and HypoM DMRs.

**Figure 2.**
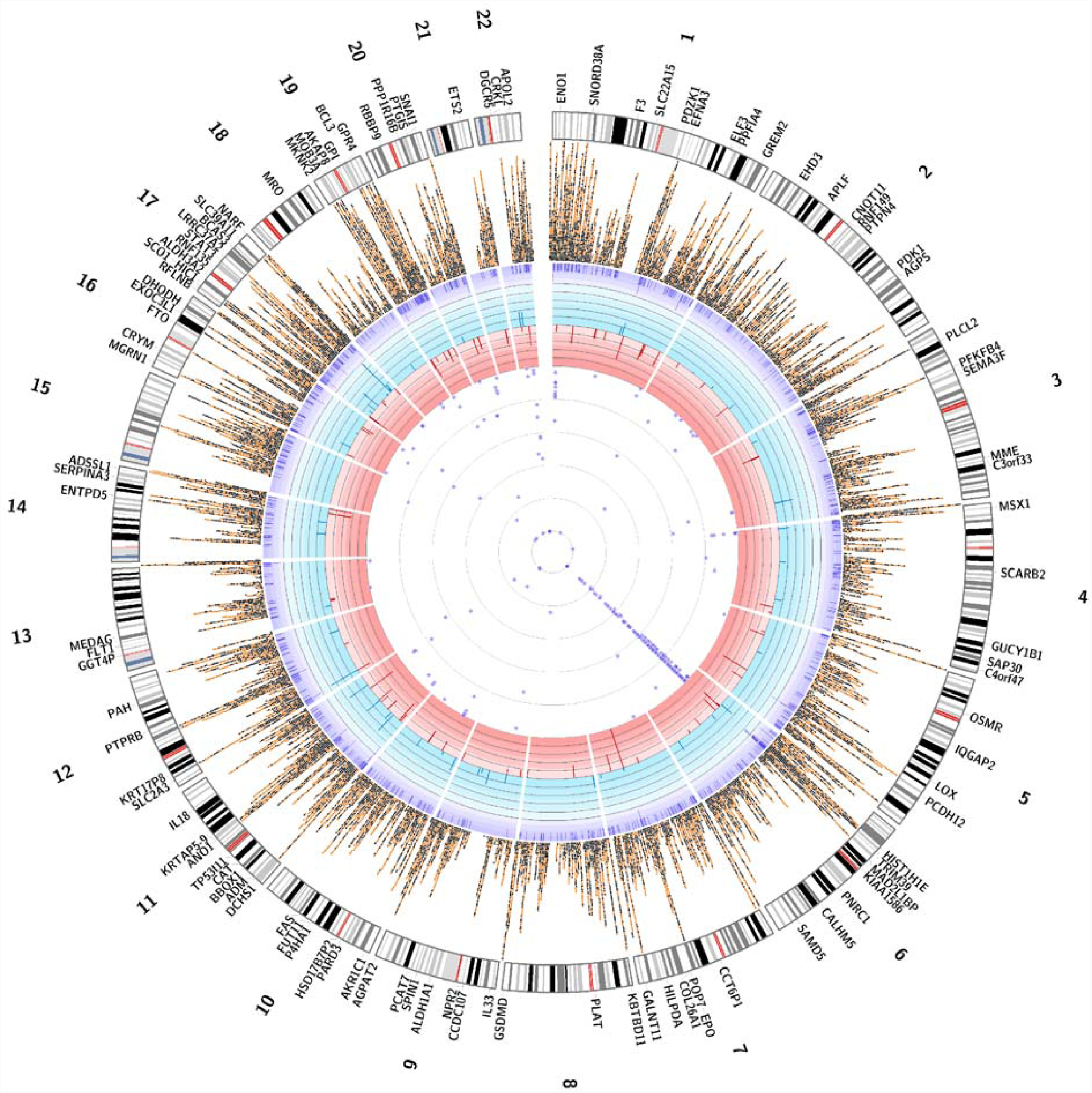
A Circos plot of significantly associated CpGs, DMRs, gene expression traits associated with WAA and GWAS catalog traits associated with replicated gene expression traits. The innermost ring represents the 257 GWAS catalog trait associated with the 28 genes replicated in the Innocenti et al. dataset (*p* < 0.05) with each purple circle represents the scaled, -log(p-value) of a study from the GWAS catalog. The second ring represents 83 negatively expressed genes associated with WAA (red bars represent fold change, ranging from 0 to −6). The third ring represents 48 positively expressed genes associated with WAA (blue bars represent fold change > 1.5, ranging from 0 to +6). The fourth ring represents 1037 DMRs significantly associated with WAA (purple tiles represents, where some DMRs are stacked when overlapping). The fifth ring represents 23,317 significantly differentially methylated CpGs (black squares represent 7913 hypomethylated CpG sites; orange circles represent 15,404 hypermethylated CpG sites; not all CpGs are depicted due to reduced crowding implemented in the Circos program). The next ring represents the karyotype of the human genome (reference hg38) and the outermost ring corresponds to the gene names of the 131 WAA-associated gene expression traits identified.

As compared to all CpG sites tested, the gene body (45.0% vs 40.9%, *p* < 0.0001, chi-square test) and shelf (8.4% vs 6.9%, *p* = 0.011, chi-square test) had significantly higher proportion of DM CpGs, while the promoter (18.4% vs 20.3%, *p* < 0.002, chi-square test), intergenic regions (IGR) (24.2% vs 27.7%, *p* < 0.0001, chi-square test) and shore regions (16.3% vs 18.2%, *p* < 0.0022, chi-square test) had significantly lower proportion of DM CpGs. In our analysis of DM loci associated with WAA, 75.9% of DM CpG sites within islands were HypoM, while the shore, shelf and open sea were predominantly HyperM (65.5%, 81.7% and 78.9% respectively) **(Supplementary Figure 2C)**. Within transcriptionally-regulated promoter regions, 54.0% of CpG sites were HypoM. Within the promoter, 71.5% of DM CpGs were HypoM 200 kb upstream of the transcription start site (TSS), while 42.9% of DM CpGs were HypoM 1500 kb upstream of the TSS. **(Supplementary Figure 2D)**. DNA methylation around TSS is an established predictor of gene expression, with increased methylation leading to decreased expression [27, 28]. In the gene body, 73.0% of DM sites were HyprM **(Supplementary Figure 2D)**. Intergenic, 5’-UTR and 3’-UTR regions were predominantly HyperM (67.5%, 64.0%, 79.6% respectively).

Next, we characterized the locations of WAA-associated CpGs by genomic features and gene annotations. Within the 7913 HypoM CpG sites associated with WAA, there was a greater proportion of CpGs located in islands. Within the 15,404 HyperM CpGs associated with WAA, there was a greater proportion in the shelf and open sea regions **(Supplementary Figure 2E)**, 5’-UTR, promoter regions and gene body **(Supplementary Figure 2F)**.

### Gene expression trait correlation to DNA methylation patterns

To determine the relationship of DMRs associated to WAA to gene expression traits, we next looked at the association of the 1034 unique genes corresponding to the 1037 DMRs with their respective gene expression profiles. Although there was no correlation of WAA-associated HyprM gene regions with gene expression (**Figure 1C**, Pearson’s r = 0.036, *p* = 0.55), HypoM gene regions associated with WAA were negatively correlated with gene expression (**Figure 1C**, Pearson’s r = −0.14, *p* = 0.009). In general, all genes within DMR associated with WAA also showed negative correlation between gene expression and methylation (Pearson’s r = −0.1, *p* = 0.013). From the 131 genes with gene expression significantly associated with WAA identified, we identified an overlap of ten DM gene regions: *COL26A1* (minFDR = 3.99 × 10^−45^, mean Beta = −0.0816, consisting of 16 CpGs), *HIC1* (minFDR = 5.35 × 10^−8^, mean Beta = 0.0011, consisting of 35 CpGs), *MKNK2* (minFDR = 8.40 × 10^−6^, mean Beta = 0.0404, consisting of 9 CpGs), *RNF135* (minFDR = 3.17 × 10^−84^, mean Beta = −0.2504, consisting of 18 CpGs) and *TRIM39* (minFDR = 4.20 × 10^−8^, mean Beta = 0.0444, consisting of 14 CpGs), with concordant directions of effect (e.g. increased methylation leading to decreased gene expression). A comprehensive Circos plot summarizes the 23,317 HypoM and HyprM CpG sites, the 1037 DMRs, the 131 genes significantly associated with WAA and the 257 GWAS studies associated with the 28 genes replicated gene expression traits found in the validation dataset of Innocenti et al. (**Figure 2, Supplemental Table 2**).

### Functional representation of WAA associated genes and DM genes associated with African ancestry

To understand the biological relevance of differentially HyprM and HypoM genes associated with African ancestry, we performed Gene Ontology (GO) analysis of the 432 unique genes comprised of the 435 HyprM DMRs and 602 unique genes comprised of the 602 HypoM DMRs using a gene panel of all annotated genes in the GO database. HyprM genes are enriched for “cell development” (BH-adjusted *p* = 0.0016) and “apoptotic process” (BH-adjusted *p* = 0.0076) within the category of biological processes **(Figure 3A)**. HypoM genes were enriched for “system development” (BH-adjusted *p* = 4.0 × 10^−6^), “response to drug” (BH-adjusted *p* = 1.7 × 10^−4^), and “response to hypoxia” (BH-adjusted *p* = 0.0068) within the category of biological processes (**Figure 3B**). In addition, HypoM genes are enriched for “sequence-specific DNA binding” (BH-adjusted *p* = 4.4 × 10^−5^) and “RNA polymerase II transcription factor activity” (BH-adjusted *p* = 4.4 × 10^−5^) within the category of molecular function **(Figure 3B)**.

**Figure 3.**
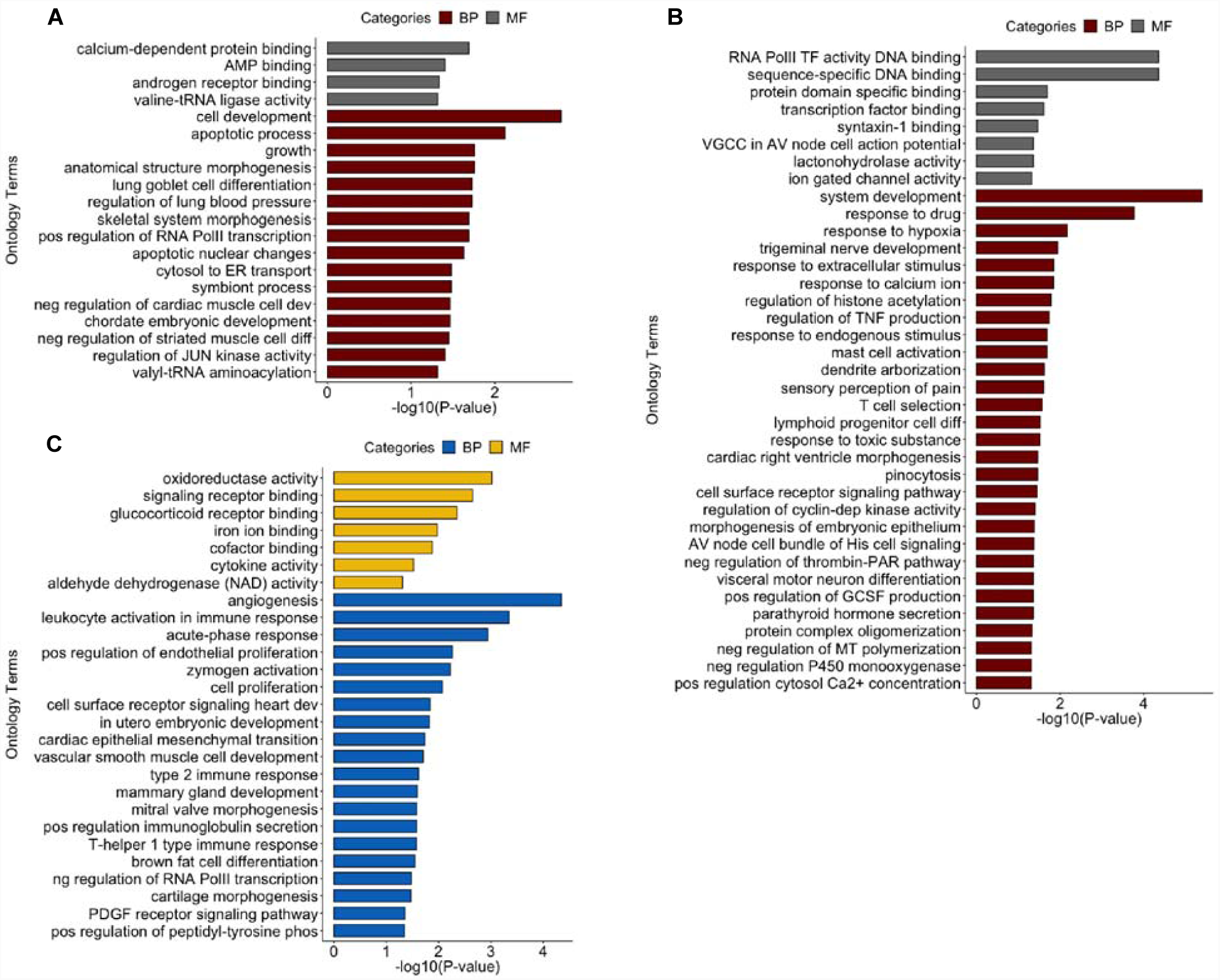
Enrichment of biological processes and molecular functions of differentially methylated genes and genes expression signatures associated with West African ancestry. Gene ontology terms that are enriched for biological processes (BP) and molecular functions (MF) for **(A)** 432 genes annotated to differentially hypermethylated regions, **(B)** 602 genes annotated to differentially hypomethylated regions and **(C)** 131 genes with gene expression traits associated to WAA (FDR < 0.10). *p*-values are BH-adjusted *p*-values obtained from gProfiler.

With respect to GO analysis of the 131 WAA-associated gene expression traits (FDR < 0.10), “angiogenesis” (BH-adjusted *p* = 4.5 × 10^−5^), “leukocyte activation involved in immune response” (BH-adjusted *p* = 4.5 × 10^−4^), “acute-phase response” (BH-adjusted *p* = 0.0011), “positive regulation of endothelial cell proliferation” (BH-adjusted *p* = 0.0055), “zymogen activation” (BH-adjusted *p* = 0.0059) and “cell proliferation” (BH-adjusted *p* = 0.0085) were biological processes that are enriched in hepatocytes **(Figure 3C)**. Molecular functions, such as “oxidoreductase activity” (BH-adjusted *p* = 9.6 × 10^−4^), “signaling receptor binding” (BH-adjusted *p* = 0.0023), “glucocorticoid receptor binding” (BH-adjusted *p* = 0.0045) and the KEGG pathway “HIF-1 signaling pathway” (BH-adjusted *p* = 8.2 × 10^−4^) were also enriched **(Figure 3C)**.

## Discussion

Several studies have shown that the first several principal components of methylation data can capture population structure in cohorts composed of European and African individuals [29]. Recently, it was shown that genetic ancestry can be used as a proxy, not only for uncovering unknown covariates contributing to epistatic and gene-environment interactions from gene expression data, but also from DNA methylation data [30]. Indeed, approximately 75% of variation in methylation was attributable to shared genomic ancestry [31]. Moreover, clinically meaningful measures can be associated to the proportion of WAA as has been shown for lung-function prediction [32].

In our study, we investigated population-specific gene expression and DNA methylation in AAs, an admixed population. We found HypoM hepatic genes, which indicate increased gene expression, are enriched for “system development” and “response to drug”. HypoM, or demethylation, may be a better predictor of gene expression than methylation which, in contrast to demethylation, may or may not affect gene expression, depending on the gene region methylated. Methylation within the TSS of the promoter is well known to repress gene expression while methylation within the gene body results in more variable expression [28, 33].

Among gene expression traits associated with WAA, we identified hepatic genes that are enriched for “angiogenesis” as the primary ontology term, followed by inflammatory response categories including “leukocyte activation in immune response”, “acute-phase response”, “positive regulation of vascular endothelial proliferation”, “zymogen activation” and “cell proliferation”. Angiogenesis and inflammatory response terms may underlie conditions that AAs may be more susceptible to, such as CVD and other chronic inflammatory disease. In particular, *APOL1, PTGIS* and *PLAT* expression, which we show to be associated with WAA, have been shown to increase CVD risk and renal disease in AAs [13, 34, 35]. Other genes associated with WAA include *ALDH1A1*, which is involved in alcohol and aldehyde metabolism disorders and cancer risk [11, 36, 37], *IL-33*, which is involved in beneficial immune response [38-40], and *VEGFA*, which has been linked to renal disease and microvascular complications of diabetes [7, 14, 41, 42].

Of the 131 genes associated with WAA, we were able to replicate a quarter within our replication and validation cohorts **(Table 2)**. We also identified an overlap with five significantly DM genes in concordant directions of effect, *COL26A1, HIC1, MKNK2, RNF135, SNAI1* and *TRIM39*. *HIC1* is a potential tumor suppressor that has been linked to poorer outcomes in laryngeal cancer in AAs [43, 44]. *RNF135* is a ring finger protein that is regulated by several population-specific variants [45]. RNF135, itself, then regulates other genes at distant loci and has been implicated in glioblastomas and autism [45-47]. Of particular interest for African ancestry populations, *RNF135* has been found to be under selective pressure specifically in African populations [48, 49].

In PharmGKB VIP genes, we identified five genes associated with WAA: *ALDH1A1, CYP2C19, P2RY1, PTGIS* and *VDR* [22, 25]. Of particular importance, we found that for every 1% increment in African ancestry, there was a corresponding 1.36% decrement in *CYP2C19* expression and 1.61% increment in *P2RY1* expression. *CYP2C19* is involved in the metabolism of many commonly prescribed drugs including clopidogrel, an antiplatelet therapy widely used for thrombo-prophylaxis of CVDs and linked to substantial inter-patient differences in drug response. It is also known to inhibit the *P2RY* family of receptor proteins on the surface of platelets [34, 35, 50, 51]. By inhibiting *P2RY12* function, clopidogrel indirectly suppresses platelet clustering and clot formation and prevents clots contributing to heart attack, stroke and deep vein thrombosis [52-54]. *P2RY1* works in concert with *P2RY12* to promote platelet activation and aggregation [55]. Consequently, *P2YR1* variants have been associated with increased platelet response to adenosine 5’-diphosphate (ADP) stimulation [56] and increased expression may be linked to thrombotic disease [57].

Clopidogrel requires *CYP2C19*-mediated conversion to its active form, but it has been shown that different populations have different levels of CYP2C19 activity [12, 58-61]. The underlying mechanism is multifactorial and genetic polymorphisms are one of the main causes of variable drug response within an individual and across populations [6, 62, 63]. A study conducted across 24 U.S. hospitals showed one-year mortality rates of 7.2% in clopidogrel-treated AAs, compared to 3.6% for Caucasians on clopidogrel [6]. This study also found that AAs were at a higher risk of cardiovascular events and mortality from poor antiplatelet response to clopidogrel. Our finding that *CYP2C19* expression is reduced with WAA while *P2RY1* expression is inversely increased, suggests that clopidogrel resistance and susceptibility of AAs to thrombotic disease may be due to ancestry-associated gene expression.

Another WAA-associated gene was *PLAT*, for which expression is decreased with increased WAA. The *PLAT* gene, which is involved in plasminogen activation and encodes tissue plasminogen activator (t-PA), is linked to thrombosis and increased risk of CVD [64, 65]. In AAs, polymorphisms in *PLAT* have been linked to CVD and higher levels of t-PA antigen seen in both myocardial infarction and venous thromboembolism [66]. In addition, increased plasma fibrinogen level, which is involved in the fibrinolytic pathway and regulated by t-PA, has been linked to increased venous thrombosis risk in AAs [67, 68].

*VDR* is also important in health and disease because Vitamin D and its active metabolite 1,25(OH)_2_D are exogenous hormones created by sun exposure or through diet, with established deficiencies in both Vitamin D and its bioactive metabolites in AAs [69, 70]. Those of African ancestry are known to have lower plasma 1,25(OH)_2_D levels and our findings support this, suggesting that this may lead to an upregulation of *VDR* with increased WAA that may compensate for these lower levels. Additionally, *VDR* single nucleotide polymorphisms (SNPs) have been implicated in CVDs and various cancer susceptibilities in AAs [8, 9, 15, 71, 72]. SNPs within *VDR* may exacerbate an already deficient vitamin D environment in AAs.

Several limitations exist in this study. First, we only had 60 primary hepatocyte cultures included in this analysis, which limits our power to detect small changes in gene expression associated with ancestry. Second, the GTEx replication liver cohort, with 15 AA livers, also severely lacked power to replicate our findings. Third, while most of the genes found in the PharmGKB VIP subset analysis were not statistically significant in the complete genome-wide set, we found *ALDH1A1* and *PTGIS* to be significantly associated to WAA in both the genome-wide and in the 64 PharmGKB VIP analysis, and we replicated 8 of the 131 significant WAA-associated genes in a GTEx liver cohort of 15 AAs. More importantly, and the reason for the low power of our dataset and any other dataset available for AAs, there are very few genome-wide datasets of both genotype and gene expression in AAs, and those that exist have similarly under-powered cohort sizes.

Another differentiation, though not considered a limitation, was that we used cultured hepatocytes as opposed to frozen liver tissues, as was the case with GTEx. Primary cultures may show differences in gene expression profiles from those seen in the intact organ, which consist of only 60% hepatocytes [73]. However, our study design has the advantage of only assaying the gene expression of a single cell type as opposed to the multiple cell types found in liver. Previous studies have shown that primary human hepatocytes show similar gene expression levels for both Phase I (CYPs) and Phase II (e.g. UGTs) drug-metabolizing enzymes to those obtained from frozen liver tissue [74, 75]. Also, the use of primary hepatocyte cultures reduces the effect of the environmental confounders inherent in liver tissue (i.e. there is less effect of previous drug/disease exposure) through the controlled tissue culture processes following hepatocyte isolation). The artifact of previous disease/drug exposures is present in all transcriptome studies conducted in post-mortem human liver tissue.

## Conclusion

Currently, genome-wide genetic, epigenetic, and multi-omics datasets of AAs are lacking in both the scientific literature and in public databases and repositories. Since genomics studies should be inclusive of all populations to comprehensively unravel disease etiology, the dearth of genetic data in minorities and underrepresented populations poses a major scientific and medical dilemma in new drug development, precision medicine, and public health policies. An archetypal example is the rs12777823 SNP in *CYP2C9*, which was found to associate with decreased warfarin dose requirement, but only in AAs [76]. Findings, such as these, were only made possible due to focused studies in a minority patient population.

In conclusion, our study has important implications in the use of genetic ancestry in understanding phenotypic differences and health disparities in AAs. Our study also has implications in determining inter-individual genetic factors and drug response outcomes in admixed populations for precision medicine. As evidenced by limited genome-wide data in AAs within public databases and biobanks, such as in GTEx, our study further stresses the need for genome-wide studies in minority and under-represented populations.

### List of Abbreviations

AA: African Americans
WAA: West African ancestry
CVD: Cardiovascular Disease
IBD: Identity-by-Descent
QC: Quality control
GWAS: Genome-wide Association Studies
eQTLs: expression quantitative trait loci
GTEx: Genotype-Tissue Expression Project
PCA: Principal Component Analysis
PEER: Probabilistic estimation of expression residuals
DE: Differentially Expressed
DMR: Differentially Expressed Region
HyprM: – Hypermethylated
HypoM: – Hypomethylated

### Declarations

#### Ethical Approval and consent to participate

The Institutional Review Board of Northwestern University has waiver the need for approval as this study used human samples obtained from deceased individuals and it thus not considered Human Subjects research.

#### Consent for publication

Consent for publication is not applicable.

#### Availability of data and materials

The dataset supporting the conclusion of this article are include with the article and is additional files. All raw gene expression and methylation array data has been submitted to the Gene Expression Omnibus (GEO accession number GSE124076) and (GEO accession number GSE123995), respectively.

#### Authors’ contributions

CSP and MAP Conception and Design, Analysis and interpretation of data, Drafting or revising the article; YX Design, Analysis and interpretation of data; TD Design and Acquisition of data; YZ Analysis; EG Analysis; ES, CA Acquisition of data.

## Acknowledgements

We would like to acknowledge Dr. Heather Wheeler for her invaluable scientific input. ERG wishes to thank the President and Fellows of Clare Hall, University of Cambridge for providing a stimulating intellectual home and for the generous support during the Easter and Lent terms.

## Supplementary Files

**Supplementary Figure 1.**
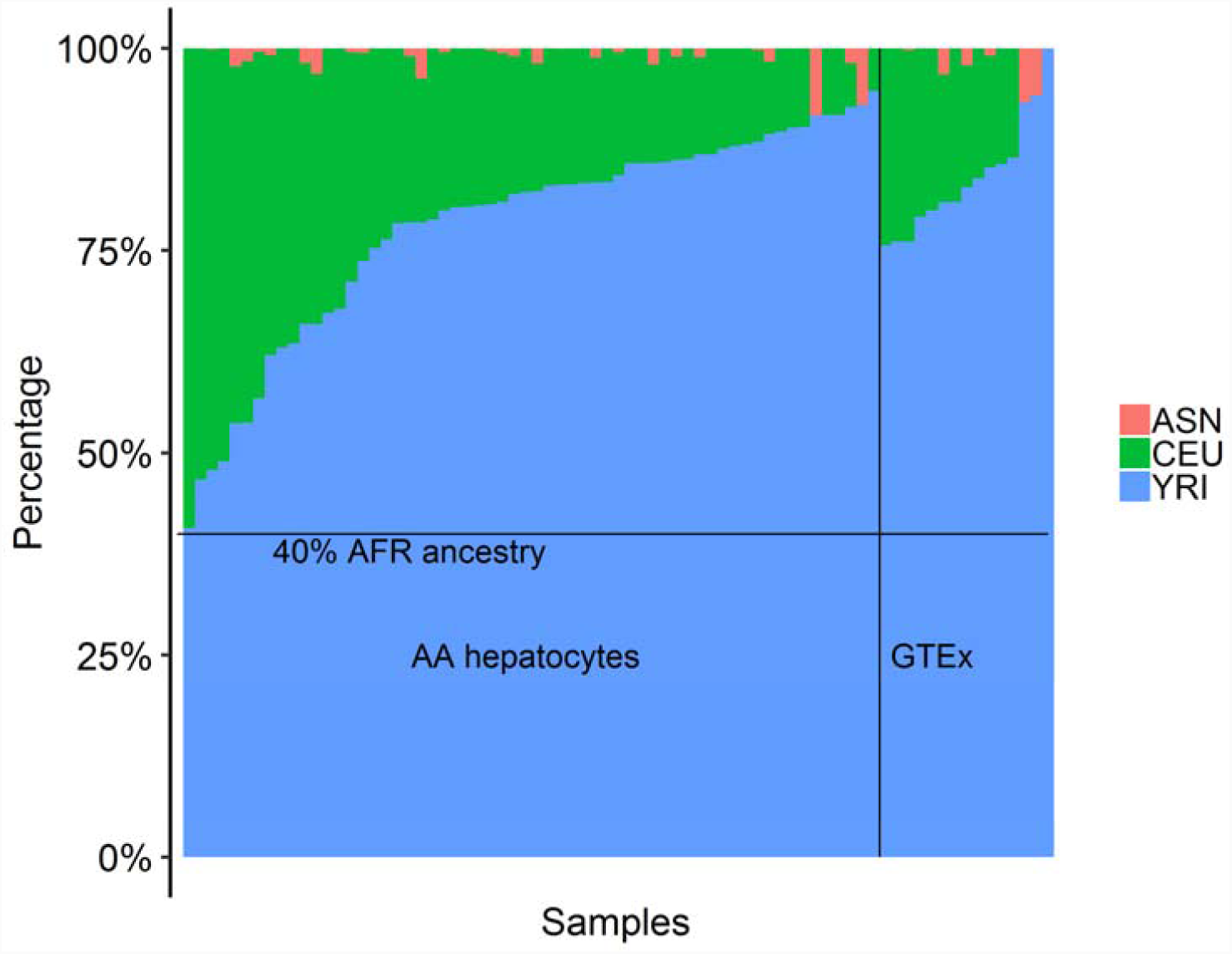
West African ancestry estimation in the AA hepatocytes and GTEx cohorts. An admixture plot showing the ancestry components of the 60 African-American hepatocyte and 15 GTEx cohorts was calculated using fastStructure v1.3. A model with three parental ancestral components (K = 3) was used to estimate from three populations from HapMap phase 3 reference data: Han Chinese in Beijing, China (CHB) and Japanese in Tokyo, Japan (JPT) combined as Asian (ASN, pink); Utah residents with Northern and Western European ancestry (CEU, green); Yoruba in Ibadan, Nigeria (YRI) as African (AFR, blue). Percent WAA is plotted on the y-axis. Each sample is represented individually on the x-axis and partitioned into segments proportional to the contributions of the ancestral components to the individual’s genome. The black line identified the threshold use for inclusion in the study.

**Supplementary Figure 2.**
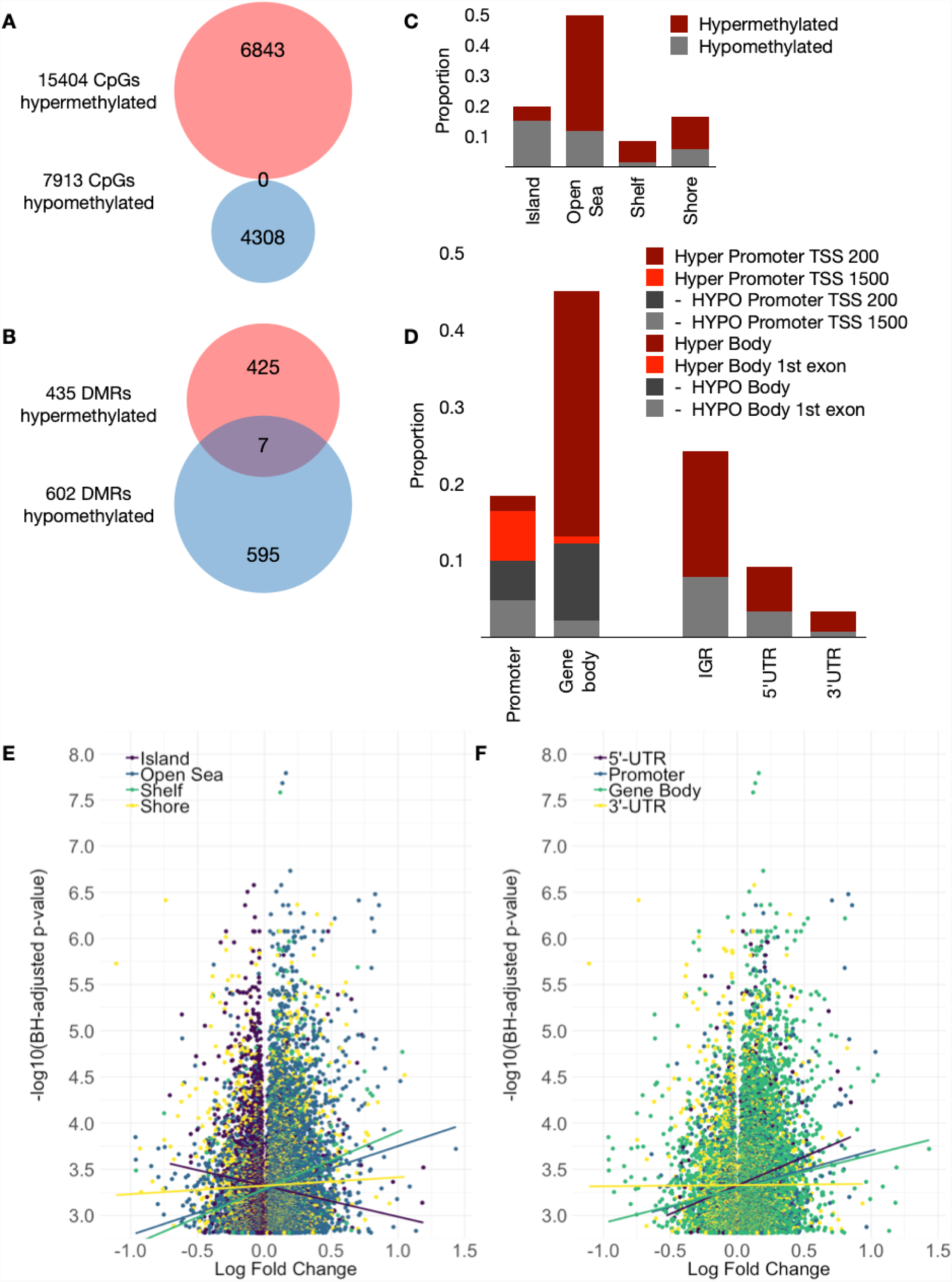
DNA methylation by genomic features and regions associated with West African ancestry. Venn diagrams of **(A)** unique genes comprised of 15,404 hyper- (pink) and 7913 hypomethylated (blue) CpG sites associated with WAA, and **(B)** unique genes comprised of 435 hyper- (pink) and 602 hypomethylated (blue) differentially methylated regions (DMRs) associated with WAA, **(C)** Bar graphs showing the proportion of hyper- and hypomethylated CpG sites by genomic features(islands, open sea, shelf and shore), and **(D)** proportion of hyper- and hypomethylated CpG sites by transcriptionally-regulated regions;promoter, gene body (with 200 or 1500 bp from the gene transcriptional start site [TSS]), intergenic region (IGR), and 5’ and 3’ untranslated regions (UTR). Volcano plots showing the correlation of log fold change in DNA methylation of the 23,317 significant DM CpG sites (BH-corrected *p* < 0.05) and -log_10_ *p*-value for the association of each CpG site to WAA by **(E)** genomic feature and by **(F)** transcriptionally-regulated regions. Colored lines indicate the relative trend by subcategory.

**Supplemental Table 1.** List of 131 West African Ancestry associated genes with p-values from the replication in DE gene analysis in a cohort of AAs and EAs or in the AA GTEx liver cohort.

|  | AA Hepatocyte Cohort |  |  | DE in AA vs EA ‡ | AA GTEx Liver Cohort § |
| --- | --- | --- | --- | --- | --- |
| *Gene/ ID* | Coefficient | *p* | FDR < 0.10 | * *p* < 0.05 | * *p* < 0.05 |
| ***IL18*** | 3.152 | 7.25E-05 | 0.0471 | 0.00000126 | 0.461 |
| ***APOL1*** | -2.848 | 3.83E-05 | 0.0380 | 0.000202 | 0.792 |
| ***VEGFA*** | -2.246 | 4.91E-05 | 0.0421 | 0.000502 | 0.032 |
| ***PARD3*** | 0.437 | 1.86E-04 | 0.0773 | 0.000696 | 0.445 |
| ***SAP30*** | -1.945 | 2.93E-04 | 0.0788 | 0.00167 | 0.844 |
| ***LRRC37A2*** | -0.306 | 6.73E-04 | 0.0988 | 0.00249 | 0.514 |
| ***ENO1*** | -1.007 | 1.89E-04 | 0.0773 | 0.00266 | 0.072 |
| ***PGK1*** | -0.929 | 5.29E-04 | 0.0882 | 0.00268 | 0.518 |
| ***DGCR5*** | 2.192 | 1.62E-04 | 0.0751 | 0.00278 | 0.144 |
| ***MRO*** | 1.131 | 2.57E-05 | 0.0380 | 0.00282 | 0.113 |
| ***GREM2*** | -2.191 | 6.86E-04 | 0.0988 | 0.00327 | 0.580 |
| ***CYP21A2*** | -2.986 | 3.17E-04 | 0.0790 | 0.00385 | 0.962 |
| ***CEBPB*** | -1.454 | 6.36E-04 | 0.0967 | 0.00445 | 0.651 |
| ***MAD2L1BP*** | -0.505 | 3.04E-04 | 0.0790 | 0.00605 | 0.179 |
| ***GPR4*** | -1.316 | 3.08E-04 | 0.0790 | 0.00706 | 0.176 |
| ***RNF149*** | -0.833 | 2.37E-04 | 0.0788 | 0.00743 | 0.288 |
| ***GPI*** | -1.453 | 4.58E-05 | 0.0411 | 0.0123 | 0.048 |
| ***SLC22A15*** | -1.271 | 6.86E-04 | 0.0988 | 0.0125 | 0.943 |
| ***HIC1*** | -1.567 | 9.64E-06 | 0.0307 | 0.013 | 0.890 |
| ***APOL2*** | -1.643 | 1.06E-04 | 0.0583 | 0.0134 | 0.273 |
| ***MME*** | 2.640 | 3.94E-04 | 0.0854 | 0.0137 | 0.384 |
| ***MKNK2*** | -1.545 | 3.44E-04 | 0.0811 | 0.0262 | 0.454 |
| ***MGRN1*** | -1.010 | 2.76E-04 | 0.0788 | 0.0313 | 0.220 |
| ***C3orf33*** | 0.506 | 6.45E-04 | 0.0973 | 0.0314 | 0.475 |
| ***PTPN4*** | 0.484 | 5.37E-04 | 0.0882 | 0.0321 | 0.541 |
| ***NPR2*** | 1.235 | 6.11E-05 | 0.0471 | 0.0444 | 0.208 |
| ***MSX1*** | 1.432 | 2.90E-04 | 0.0788 | 0.0448 | 0.845 |
| ***PDK1*** | -2.038 | 6.30E-04 | 0.0966 | 0.0498 | 0.389 |
| ***ADSSL1*** | -3.254 | 3.81E-04 | 0.0846 | 0.0553 | 0.962 |
| ***STAT3*** | -0.859 | 2.99E-04 | 0.0790 | 0.0561 | 0.366 |
| ***EXOC3L1*** | -1.518 | 4.67E-04 | 0.0868 | 0.0599 | 0.859 |
| ***ANO1*** | 2.336 | 2.41E-04 | 0.0788 | 0.0637 | 0.372 |
| ***CCT6P1*** | -0.765 | 5.09E-04 | 0.0873 | 0.0642 | 0.345 |
| *SERPINA3* | -3.379 | 2.79E-04 | 0.0788 | 0.071 | 0.493 |
| *AGPS* | 0.894 | 5.31E-04 | 0.0882 | 0.0796 | 0.692 |
| *ALDH1A1* | 2.077 | 6.56E-05 | 0.0471 | 0.0823 | 0.558 |
| *PLAT* | -1.883 | 3.49E-05 | 0.0380 | 0.0984 | 0.823 |
| *AKR1C1* | 2.026 | 5.38E-04 | 0.0882 | 0.102 | 0.397 |
| *IL33* | -1.069 | 6.76E-05 | 0.0471 | 0.103 | 0.922 |
| *CAT* | 1.183 | 2.88E-04 | 0.0788 | 0.117 | 0.664 |
| *IQGAP2* | 0.805 | 2.68E-04 | 0.0788 | 0.118 | 0.222 |
| *TRIM39* | -0.606 | 4.89E-04 | 0.0868 | 0.119 | 0.030 |
| *LOX* | -4.029 | 2.81E-05 | 0.0380 | 0.125 | 0.860 |
| *MEDAG* | -0.732 | 2.63E-04 | 0.0788 | 0.128 | 0.249 |
| *PCAT7* | 1.141 | 4.04E-04 | 0.0856 | 0.128 | NT |
| *MOB3A* | -1.408 | 2.62E-04 | 0.0788 | 0.135 | 0.933 |
| *SCARB2* | 0.883 | 4.64E-04 | 0.0868 | 0.138 | 0.289 |
| *POP7* | -1.209 | 5.54E-04 | 0.0901 | 0.139 | 0.322 |
| *PTPRB* | -0.725 | 2.73E-04 | 0.0788 | 0.151 | 0.227 |
| *OSMR* | -2.193 | 9.69E-06 | 0.0307 | 0.156 | 0.856 |
| *SLC39A11* | 0.491 | 3.02E-05 | 0.0380 | 0.159 | 0.404 |
| *HILPDA* | -2.583 | 9.83E-06 | 0.0307 | 0.171 | 0.203 |
| *NOVA2* | -0.479 | 6.10E-04 | 0.0942 | 0.174 | 0.186 |
| *BBOX1* | 3.049 | 8.23E-05 | 0.0485 | 0.195 | 0.843 |
| *EFNA3* | -1.606 | 1.63E-04 | 0.0751 | 0.203 | 0.314 |
| *ENTPD5* | 1.265 | 4.65E-04 | 0.0868 | 0.222 | 0.744 |
| *F3* | -1.373 | 5.65E-04 | 0.0901 | 0.231 | 0.599 |
| *GSDMD* | -1.104 | 3.50E-04 | 0.0814 | 0.235 | 0.502 |
| *PTGIS* | -0.855 | 4.24E-06 | 0.0307 | 0.237 | 0.808 |
| *ADM* | -3.147 | 1.71E-04 | 0.0770 | 0.241 | 0.430 |
| *CRYM* | 1.573 | 3.69E-05 | 0.0380 | 0.249 | 0.213 |
| *PPP1R16B* | -0.553 | 4.33E-04 | 0.0868 | 0.249 | 0.979 |
| *SLC2A3* | -1.961 | 2.99E-05 | 0.0380 | 0.255 | 0.032 |
| *GALNT11* | 0.615 | 1.79E-04 | 0.0773 | 0.258 | 0.767 |
| *C4orf47* | -0.483 | 3.41E-04 | 0.0811 | 0.264 | 0.069 |
| *ALDH3A2* | 1.196 | 3.93E-04 | 0.0854 | 0.265 | 0.765 |
| *PFKFB4* | -2.528 | 1.44E-04 | 0.0733 | 0.266 | 0.289 |
| *EPO* | -6.104 | 4.75E-04 | 0.0868 | 0.272 | 0.116 |
| *FAS* | 1.317 | 4.94E-04 | 0.0868 | 0.277 | 0.422 |
| *DCHS1* | -1.391 | 5.02E-04 | 0.0868 | 0.28 | 0.198 |
| *ETS2* | -2.109 | 4.33E-04 | 0.0868 | 0.282 | 0.597 |
| *KIAA1586* | 0.919 | 3.25E-04 | 0.0796 | 0.314 | 0.470 |
| *RNF135* | 1.801 | 2.67E-05 | 0.0380 | 0.318 | 0.620 |
| *COL26A1* | 0.448 | 4.88E-04 | 0.0868 | 0.325 | 0.037 |
| *PCDH12* | -1.693 | 5.35E-04 | 0.0882 | 0.328 | 0.857 |
| *P4HA1* | -2.038 | 2.29E-05 | 0.0380 | 0.377 | 0.717 |
| *SAMD5* | 1.536 | 3.67E-04 | 0.0823 | 0.396 | 0.973 |
| *TP53I11* | -1.686 | 4.29E-04 | 0.0868 | 0.401 | 0.235 |
| *PAH* | 1.446 | 2.76E-04 | 0.0788 | 0.43 | 0.149 |
| *NARF* | -1.272 | 1.57E-04 | 0.0751 | 0.433 | 0.149 |
| *CRKL* | -0.696 | 2.80E-04 | 0.0788 | 0.438 | 0.795 |
| *LRG1* | -2.859 | 2.52E-04 | 0.0788 | 0.444 | 0.949 |
| *SIVA1* | -2.137 | 5.42E-06 | 0.0307 | 0.449 | 0.361 |
| *FUT11* | -1.749 | 7.09E-05 | 0.0471 | 0.455 | 0.550 |
| *BCL3* | -1.860 | 4.01E-04 | 0.0856 | 0.474 | 0.966 |
| *EFNA1* | -3.540 | 7.23E-05 | 0.0471 | 0.476 | 0.970 |
| *AKAP8* | -0.812 | 1.07E-05 | 0.0307 | 0.48 | 0.764 |
| *SPIN1* | 0.675 | 1.52E-05 | 0.0358 | 0.49 | 0.678 |
| *HSD17B7P2* | -1.548 | 1.96E-04 | 0.0785 | 0.491 | 0.027 |
| *SEMA3F* | -1.230 | 5.74E-04 | 0.0901 | 0.532 | 0.311 |
| *SNAI1* | -1.791 | 1.11E-04 | 0.0583 | 0.55 | 0.360 |
| *BCAS3* | 0.178 | 4.95E-04 | 0.0868 | 0.556 | 0.994 |
| *RBBP9* | 0.911 | 4.12E-04 | 0.0862 | 0.562 | 0.473 |
| *PLCL2* | 0.760 | 6.32E-05 | 0.0471 | 0.568 | 0.046 |
| *EHD3* | -0.769 | 1.89E-04 | 0.0773 | 0.572 | 0.720 |
| *SCO1* | 0.528 | 4.77E-04 | 0.0868 | 0.583 | 0.150 |
| *ELF3* | -2.508 | 2.48E-04 | 0.0788 | 0.59 | 0.994 |
| *PNRC1* | -1.242 | 5.69E-04 | 0.0901 | 0.599 | 0.805 |
| *CCDC107* | -0.787 | 4.63E-04 | 0.0868 | 0.637 | 0.247 |
| *FTO* | 0.280 | 6.79E-04 | 0.0988 | 0.651 | 0.120 |
| *CNOT11* | -0.915 | 1.11E-04 | 0.0583 | 0.672 | 0.409 |
| *DHODH* | -1.893 | 2.38E-04 | 0.0788 | 0.678 | 0.048 |
| *AGPAT2* | -1.399 | 3.13E-04 | 0.0790 | 0.68 | 0.962 |
| *KBTBD11* | 1.744 | 2.83E-04 | 0.0788 | 0.696 | 0.083 |
| *KRTAP5-9* | -2.298 | 2.33E-04 | 0.0788 | 0.746 | 0.691 |
| *PDZK1* | 1.123 | 4.64E-04 | 0.0868 | 0.755 | 0.260 |
| *HIST1H1E* | 1.981 | 4.98E-04 | 0.0868 | 0.792 | 0.133 |
| *APLF* | 0.153 | 4.79E-04 | 0.0868 | 0.814 | 0.367 |
| *CDK18* | -1.805 | 2.48E-04 | 0.0788 | 0.89 | 0.686 |
| *FLT1* | -0.726 | 3.60E-04 | 0.0823 | 0.918 | 0.952 |
| *CALHM5* | -0.457 | 6.58E-04 | 0.0985 | NT | 0.486 |
| *GGT1* | 2.155 | 8.14E-05 | 0.0485 | NT | 0.888 |
| *GGT4P* | 0.914 | 2.32E-04 | 0.0788 | NT | NT |
| *GUCY1B1* | 1.015 | 3.43E-04 | 0.0811 | NT | 0.743 |
| *KRT17P8* | 4.401 | 2.80E-04 | 0.0788 | NT | 0.647 |
| *PPFIA4* | -2.067 | 2.56E-04 | 0.0788 | NT | 0.509 |
| *RFLNB* | -3.175 | 1.14E-05 | 0.0307 | NT | 0.275 |
| *SNORD38A* | -2.588 | 6.74E-04 | 0.0988 | NT | NT |
| *ENSG00000173727* | -1.267 | 9.39E-05 | 0.0536 | NT | 0.910 |
| *ENSG00000205622* | -1.547 | 2.04E-05 | 0.0380 | NT | 0.174 |
| *ENSG00000224093* | -1.386 | 3.65E-04 | 0.0823 | NT | 0.324 |
| *ENSG00000250764* | -0.540 | 3.53E-05 | 0.0380 | NT | 0.944 |
| *ENSG00000258377* | 1.180 | 3.18E-04 | 0.0790 | NT | NT |
| *ENSG00000259357* | 1.869 | 8.06E-05 | 0.0485 | NT | NT |
| *ENSG00000263843* | 1.158 | 5.99E-04 | 0.0933 | NT | 0.129 |
| *ENSG00000264456* | 2.707 | 1.50E-04 | 0.0744 | NT | 0.747 |
| *ENSG00000268230* | -1.522 | 4.88E-04 | 0.0868 | NT | NT |
| *ENSG00000271239* | 5.823 | 4.16E-05 | 0.0392 | NT | 0.339 |
| *ENSG00000273259* | -2.002 | 5.71E-04 | 0.0901 | NT | NT |
| *ENSG00000275285* | -2.328 | 4.17E-04 | 0.0863 | NT | NT |
| *ENSG00000281655* | -4.148 | 2.19E-04 | 0.0788 | NT | NT |

\* Replication *p*-value based on 131 discovery genes in 60 AA Hepatocyte
‡ Replication in differentially expressed (DE) genes between 23 African-Americans (AAs) and 183 European-Americans (EAs) [23]
§ Replication of genes expressed in 15 AA in GTEx liver cohort with greater than 40% West African ancestry
NT – Not tested - were not present on the gene expression microarray
**Bolded** gene names indicated those with $p < 0.05$ in the Replication Cohort. Genes in **shaded**
rows were replicated in GTEx

**Supplemental Table 2. Phenotypic traits associated with replicated genes in the GWAS Catalog**

Refer to Excel datasheet: Supp_Table_2_gwas_catalog.xls

